# Fecal short-chain fatty acids are not predictive of colonic tumor status and cannot be predicted based on bacterial community structure

**DOI:** 10.1101/604678

**Authors:** Marc A. Sze, Begüm D. Topçuoğlu, Nicholas A. Lesniak, Mack T. Ruffin, Patrick D. Schloss

## Abstract

Colonic bacterial populations are thought to have a role in the development of colorectal cancer with some protecting against inflammation and others exacerbating inflammation. Short-chain fatty acids (SCFAs) have been shown to have anti-inflammatory properties and are produced in large quantities by colonic bacteria which produce SCFAs by fermenting fiber. We assessed whether there was an association between fecal SCFA concentrations and the presence of colonic adenomas or carcinomas in a cohort of individuals using 16S rRNA gene and metagenomic shotgun sequence data. We measured the fecal concentrations of acetate, propionate, and butyrate within the cohort and found that there were no significant associations between SCFA concentration and tumor status. When we incorporated these concentrations into random forest classification models trained to differentiate between people with normal colons and those with adenomas or carcinomas, we found that they did not significantly improve the ability of 16S rRNA gene or metagenomic gene sequence-based models to classify individuals. Finally, we generated random forest regression models trained to predict the concentration of each SCFA based on 16S rRNA gene or metagenomic gene sequence data from the same samples. These models performed poorly and were able to explain at most 14% of the observed variation in the SCFA concentrations. These results support the broader epidemiological data that questions the value of fiber consumption for reducing the risks of colorectal cancer. Although other bacterial metabolites may serve as biomarkers to detect adenomas or carcinomas, fecal SCFA concentrations have limited predictive power.

**Importance:** Considering colorectal cancer is the third leading cancer-related cause of death within the United States, it is important to detect colorectal tumors early and to prevent the formation of tumors. Short-chain fatty acids (SCFAs) are often used as a surrogate for measuring gut health and for being anti-carcinogenic because of their anti-inflammatory properties. We evaluated the fecal SCFA concentration of a cohort of individuals with varying colonic tumor burden who were previously analyzed to identify microbiome-based biomarkers of tumors. We were unable to find an association between SCFA concentration and tumor burden or use SCFAs to improve our microbiome-based models of classifying people based on their tumor status. Furthermore, we were unable to find an association between the fecal community structure and SCFA concentrations. Our results indicate that the association between fecal SCFAs, the gut microbiome, and tumor burden is weak.

---

Colorectal cancer is the third leading cancer-related cause of death within the United States (1). Less than 10% of cases can be attributed to genetic risk factors (2). This leaves a significant role for environmental, behavioral, and dietary factors (3, 4). Colorectal cancer is thought to be initiated by a series of mutations that accumulate as the mutated cells begin to proliferate leading to adenomatous lesions, which are succeeded by carcinomas (2). Throughout this progression, there are ample opportunities for bacterial populations to have a role as some bacteria are known to cause mutations, induce inflammation, and accelerate tumorigenesis (5–7). Additional cross sectional studies in humans have identified microbiome-based biomarkers of disease (8). These studies suggest that in some cases, it is the loss of bacterial populations that produce short-chain fatty acids (SCFAs) that results in increased inflammation and tumorigenesis.

Many microbiome studies use the concentrations of SCFAs and the presence of 16S rRNA gene sequences from organisms and the genes involved in producing them as a biomarker of a healthy microbiota (9, 10). Depending on the concentrations, SCFAs can have proliferative activities at low concentrations or anti-proliferative activities at higher concentrations; they can also have anti-inflammatory activities (11). Direct supplementation of SCFAs or feeding of fiber caused an overall reduction in tumor burden in mouse models of colorectal cancer (12). These results suggest that supplementation with fiber, which many colonic bacteria ferment to produce SCFAs, may confer beneficial effects against colorectal cancer. Regardless, there is a lack of consistent evidence that increasing SCFA concentrations can protect against colorectal cancer in humans. Case-control studies that have investigated possible associations between SCFAs and colon tumor status have been plagued by relatively small numbers of subjects, but have reported increased total and relative fecal acetate levels and decreased relative fecal butyrate concentrations in subjects with colonic lesions (13). In randomized controlled trials fiber supplementation has been inconsistently associated with protection against tumor formation and recurrence (14, 15). Such studies are confounded by difficulties ensuring subjects took the proper dose and using subjects with prior polyp history who may be beyond a point of benefiting from fiber supplementation. Together, these findings temper enthusiasm for treatments that target the production of SCFAs or for using them as biomarkers for protection against tumorigenesis.

## Fecal SCFA concentrations did not vary with diagnosis or treatment

To test for a significant association between colorectal cancer and SCFAs, we quantified the concentration of acetate, propionate, and butyrate in feces of previously characterized individuals with normal colons (N=172) and those with colonic adenomas (N=198) or carcinomas (N=120) (16). We were unable to detect a significant difference in any SCFA concentration across the diagnoses groups (all P>0.15; Figure 1A). Among the individuals with adenomas and carcinomas, a subset (N_adenoma_=41, N_carcinoma_=26) were treated and sampled a year later (17). None of the individuals showed signs of recurrence and yet none of the SCFAs exhibited a significant change with treatment (all P>0.058; Figure 1B). For both the pre-treatment cross-sectional data and the pre/post treatment data, we also failed to detect any significant differences in the relative concentrations of any SCFAs (P>0.16). Finally, we pooled the SCFA concentrations on a total and per molecule of carbon basis and again failed to observe any significant differences (P>0.077). Although some of the P-values from our analyses were close to 0.05, the effect sizes were all relatively small and inconsistent given the disease progression (Figure 1). These results demonstrated that there were no significant associations between fecal SCFA concentration and diagnosis or treatment.

**Figure 1.**
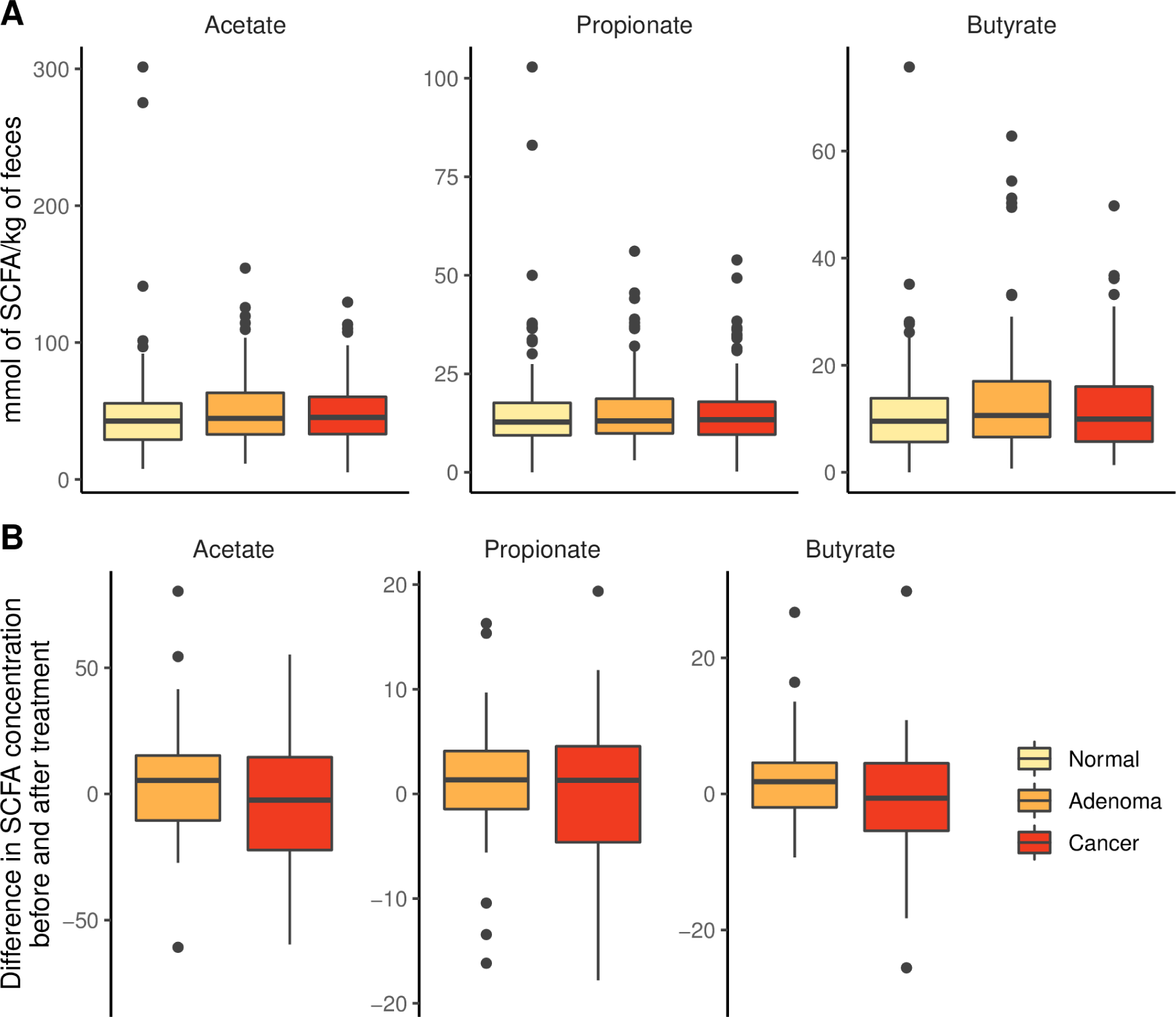
SCFA concentrations did not vary meaningfully with diagnosis of colonic lesions or with treatment for adenomas or carcinomas. (A) The concentration of fecal SCFAs from individuals with normal colons (N=172) or those with adenoma (N=198) or carcinomas (N=120). (B) A subset of individuals diagnosed with adenomas (N=41) or carcinomas (N=26) who underwent treatment were resampled a year after the initial sampling; one extreme propionate value (124.4 mmol/kg) was included in the adenoma analysis but censored from the visualization for clarity.

## Combining SCFA and microbiome data does not improve the ability to diagnose individual as having adenomas or carcinomas using a random forest model

We previously found that binning 16S rRNA gene sequence data into operational taxonomic units (OTUs) based on 97% similarity or into genera enabled us to classify individuals as having adenomas or carcinomas using random forest machine learning models (8, 16). We repeated that analysis but added the concentration of the SCFAs as possible features to train the models (Figure S1). Models trained using SCFAs to classify individuals as having adenomas or carcinomas rather than normal colons had median areas under the receiver operator characteristic curve (AUROC) that were significantly greater than 0.5 (P_adenoma_<0.001 and P_carcinoma_<0.001). However, the AUROC values to detect the presence of adenomas or carcinomas were only 0.54 and 0.55, respectively, indicating that SCFAs had poor predictive power on their own (Figure 2A). When we trained the models with the SCFAs concentrations and OTU or genus-level relative abundances the AUROC values were not significantly different from the same models trained without the SCFA concentrations (P>0.15; Figure 2A). These data demonstrate that knowledge of the SCFA profile from a subject’s fecal sample did not improve the ability to diagnose a colonic lesion.

**Figure 2.**
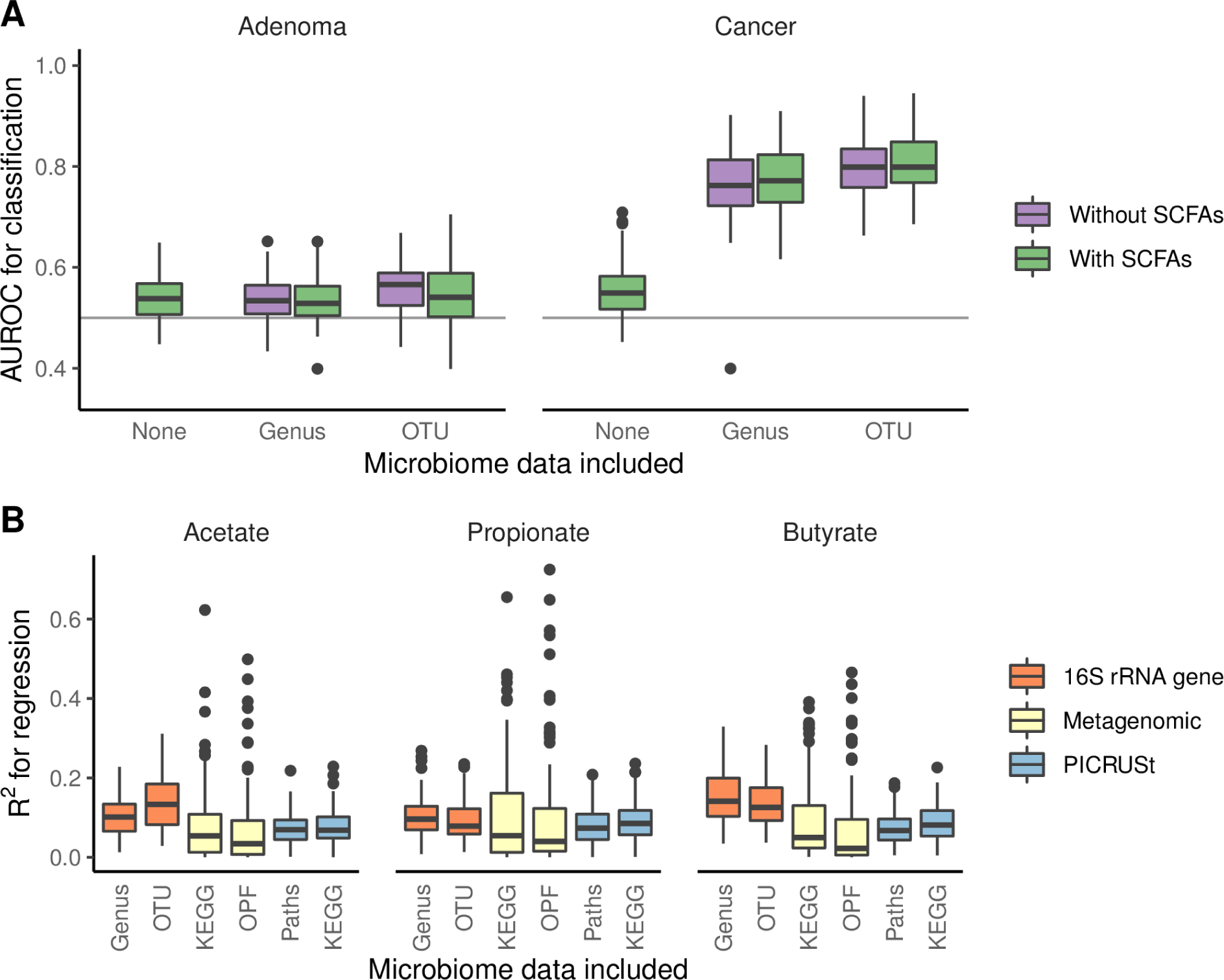
SCFA concentrations do not improve models for diagnosing the presence of adenomas, carcinomas, or all lesions and cannot be reliably predicted from 16S rRNA gene or metagenomic sequence data. (A) The median AUROC for diagnosing individuals as having adenomas or carcinomas using SCFAs was slightly better than than chance (depicted by horizontal line at 0.50), but did not improve performance of the models generated using 16S rRNA gene sequence data. (B) Regression models that were trained using 16S rRNA gene sequence, metagenomic, and PICRUSt data to predict the concentrations of SCFAs performed poorly (all median R^2^ values < 0.14). Regression models generated using 16S rRNA gene sequence and PICRUSt data included data from 490 samples and those generated using metagenomic data included data from 78 samples.

## Knowledge of microbial community structure does not predict SCFA concentrations using a random forest model

We next asked whether the fecal community structure was predictive of fecal SCFA concentrations, regardless of a person’s diagnosis. We trained random forest regression models using 16S rRNA gene sequence data binned into OTUs and genera to predict the concentration of the SCFAs (Figure S2). The largest R^2^ between the observed SCFA concentrations and the modeled concentrations was 0.14, which was observed when using genus data to predict butyrate concentrations (Figure 2B). We also used a smaller dataset of shotgun metagenomic sequencing data generated from a subset of our cohort (N_normal_=27, N_adenoma_=25, and N_cancer_=26) (18). We binned genes extracted from the assembled metagenomes into operational protein families (OPFs) or KEGG categories and trained random forest regression models using metagenomic sequence data to predict the concentration of the SCFAs (Figure S2). Similar to the analysis using 16S rRNA gene sequence data, the metagenomic data was not predictive of SCFA concentration. The largest R^2^ was 0.055, which was observed when using KEGG data to predict propionate concentrations (Figure 2B). Because of the limited number of samples that we were able to generate metagenomic sequence data from, we used our 16S rRNA gene sequence data to impute metagenomes that were binned into metabolic pathways or KEGG categories using PICRUSt (Figure S2). SCFA concentrations could not be predicted based on the imputed metagenomic data. The largest R^2^ was 0.085, which was observed when using KEGG data to predict propionate concentrations (Figure 2B). The inability to model SCFA concentrations from microbiome data indicates that the knowledge of the abundance of organisms and their genes was insufficient to predict fecal SCFA concentrations.

## Conclusion

Our data indicate that fecal SCFA concentrations are not associated with the presence of adenomas or carcinomas and that they provide weak predictive power to improve the ability to diagnose someone with one of these lesions. Furthermore, knowledge of the taxonomic and genetic structure of gut microbiota was not meaningfully predictive of SCFA concentrations. These results complement existing literature that suggest that fiber consumption and the production of SCFAs are unable to prevent the risk of developing colonic tumors. It is important to note that our analysis was based on characterizations of SCFA and microbiome profiles using fecal samples at a single time point. Furthermore, observations along the mucosa near the site of lesions may provide a stronger association. This may be a cautionary result to temper enthusiasm for SCFAs as a biomarker of gut health more generally. Going forward it is critical to develop additional hypotheses for how the microbiome and host interact to drive tumorigenesis so that we can better understand tumorigenesis and identify biomarkers that will allow early detection of lesions.

## Acknowledgements

The authors thank the Great Lakes-New England Early Detection Research Network for providing the fecal samples that were used in this study. We would thank the University of Michigan Center for Microbial Systems for enabling our short-chain fatty acid analysis. Support for MAS came from the Canadian Institute of Health Research and the National Institutes of Health (UL1TR002240). This work was also supported by the National Institutes of Health (P30DK034933 and R01CA215574).

## Materials and Methods

### Study design and sampling

The overall study design and the resulting sequence data have been previously described (16, 17). In brief, fecal samples were obtained from 172 individuals with normal colons, 198 individuals with colonic adenomas, and 120 individuals with carcinomas. Of the individuals diagnosed as having adenomas or carcinomas, a subset (N_adenoma_=41 and N_carcinoma_=26) were sampled after treatment of the lesion (median=255 days between sampling, IQR=233 to 334 days). Tumor diagnosis was made by colonoscopic examination and histopathological review of the biopsies (16). The University of Michigan Institutional Review Board approved the studies that generated the samples and informed consent was obtained from all participants in accordance to the guidelines set out by the Helsinki Declaration.

### Measuring specific SCFAs

The measurement of acetate, propionate, isobutyrate, and butyrate used a previously published protocol that used High-Performance Liquid Chromatography (HPLC) (19). Two changes were made to the protocol. First, instead of using fecal samples suspended in DNA Genotek OmniGut tubes, we suspended frozen fecal samples in 1 mL of PBS. Second, instead of using the average weight of fecal sample aliquots to normalize SCFA concentrations, we used the actual weight of the fecal samples. These methodological changes did not affect the range of concentrations of these SCFAs between the two studies. The concentrations of isobutyrate were consistently at or below the limit of detection and were not included in our analysis.

### 16S rRNA gene sequence data analysis

Sequence data from Baxter et al. (16) and Sze et al. (17) were obtained from the Sequence Read Archive (studies SRP062005 and SRP096978) and reprocessed using mothur v.1.42 (20). The original studies generated sequence data from V4 region of the 16S rRNA gene using paired 250 nt reads on an Illumina MiSeq sequencer. The resulting sequence data were assembled into contigs and screened to remove low quality contigs and chimeras. The curated sequences were then clustered into OTUs at a 97% similarity threshold and assigned to the closest possible genus with an 80% confidence threshold trained on the reference collection from the Ribosomal Database Project (v.16). We used PICRUSt (v.2.1.0-b) with the recommended standard operating protocol to generate imputed metagenomes based on the expected metabolic pathways and KEGG categories (21).

### Metagenomic DNA sequence analysis

A subset of the samples from the samples described by Baxter et al. (16) were used to generate metagenomic sequence data (N_normal_=27, N_adenoma_=25, and N_cancer_=26). These data were generated by Hannigan et al. (18) and deposited into the Sequence Read Archive (study SRP108915). Fecal DNA was subjected to shotgun sequencing on an Illumina HiSeq using 125 bp paired end reads. The archived sequences were already quality filtered and aligned to the human genome to remove contaminating sequence data. We downloaded the sequences and assembled them into contigs using MEGAHIT (22), which were used to identify open reading frames (ORFs) using Prodigal (23). We determined the abundance of each ORF by mapping the raw reads back to the ORFs using Diamond (24). We clustered the ORFs into operational protein families (OPFs) in which the clustered ORFs were more than 40% identical to each other using mmseq2 (25). We also used mmseq2 to map the ORFs to the KEGG database and clustered the ORFs according to which category the ORFs mapped.

### Random forest models

The classification models were built to predict lesion type from microbiome information with or without SCFA concentrations. The regression models were built to predict the SCFA concentrations of acetate, butyrate, and propionate from microbiome information. For classification and regression models, we pre-processed the features by scaling them to vary between zero and one. Features with no variance in the training set were removed from both the training and testing sets. We randomly split the data into training and test sets so that the training set consisted of 80% of the full dataset while the test set was composed of the remaining data. The training set was used for hyperparameter selection and training the model and the test set was used for evaluating prediction performance. For each model, the best performing hyperparameter, mtry, was selected in an internal five-fold cross-validation of the training set with 100 randomizations. The mtry parameter represents the number of features randomly sampled from the available features at a question point in the classification tree (i.e. called splits of nodes) that, when answered, lead to the greatest improvement in classification. Six values of mtry were tested and the value that provided the largest AUROC or R^2^ was selected. We trained the random forest model using the selected mtry value and predicted the held-out test set. The data-split, hyperparameter selection, training and testing steps were repeated 100 times to get a reliable and robust reading of model prediction performance. We used AUROC and R^2^ as the prediction performance metric for classification and regression models, respectively. We used the randomForest R package (version 4.6-14) as implemented in the caret R package (version 6.0-81) for developing and testing our models.

### Statistical analysis workflow

Data summaries, statistical analysis, and data visualizations were performed using R (v.3.5.1) with the tidyverse package (v.1.2.1). To assess differences in SCFA concentrations between individuals normal colons and those with adenomas or carcinomas, we used the Kruskal-Wallis rank sum test. If a test had a P-value below 0.05, we then applied a pairwise Wilcoxon rank sum test with a Benjamini-Hochberg correction for multiple comparisons. To assess differences in SCFA concentrations between individuals samples before and after treatment we used paired Wilcoxon rank sum tests to test for significance. To compare the median AUCROC for the held out data for the model generated using only the SCFAs, we compared the distribution of the data to the expected median of 0.5 using the Wilcoxon rank sum test to test whether the model performed better than would be achieved by randomly assigning the data to each diagnosis. When we compared the random forest models generated without and with SCFA data included, we used Wilcoxon rank sum tests to determine whether the models with the SCFA data included did better.

### Code availability

The code for all sequence curation and analysis steps including an Rmarkdown version of this manuscript is available at https://github.com/SchlossLab/Sze_SCFACRC_mBio_2019/.

---

**Figure S1. Comparison of training and testing results for classification models shows that the models are robust and are not overfit.** random forest classification models were generated to differentiate between individuals with normal colons and those with adenomas or carcinomas using 16S rRNA gene sequence data that were clustered into genera or OTUs with and without including the three SCFAs as additional features. random forest classification models were generated by partitioning the samples into a training set with 80% of the data and a testing set with the remaining samples for 100 randomizations.

**Figure S2. Comparison of training and testing results for regression models shows that the models are robust and are not overfit.** random forest regression models were generated to predict the concentration of each SCFA using each individuals’ microbiome data generated using 16S rRNA gene sequence and metagenomic sequence data. These regression models were generated by partitioning the samples into a training set with 80% of the data and a testing set with the remaining samples for 100 randomizations.

