## Supplementary figures and images for "Fecal short-chain fatty acids are not predictive of colonic tumor status and cannot be predicted based on bacterial community structure"

### Figure S1

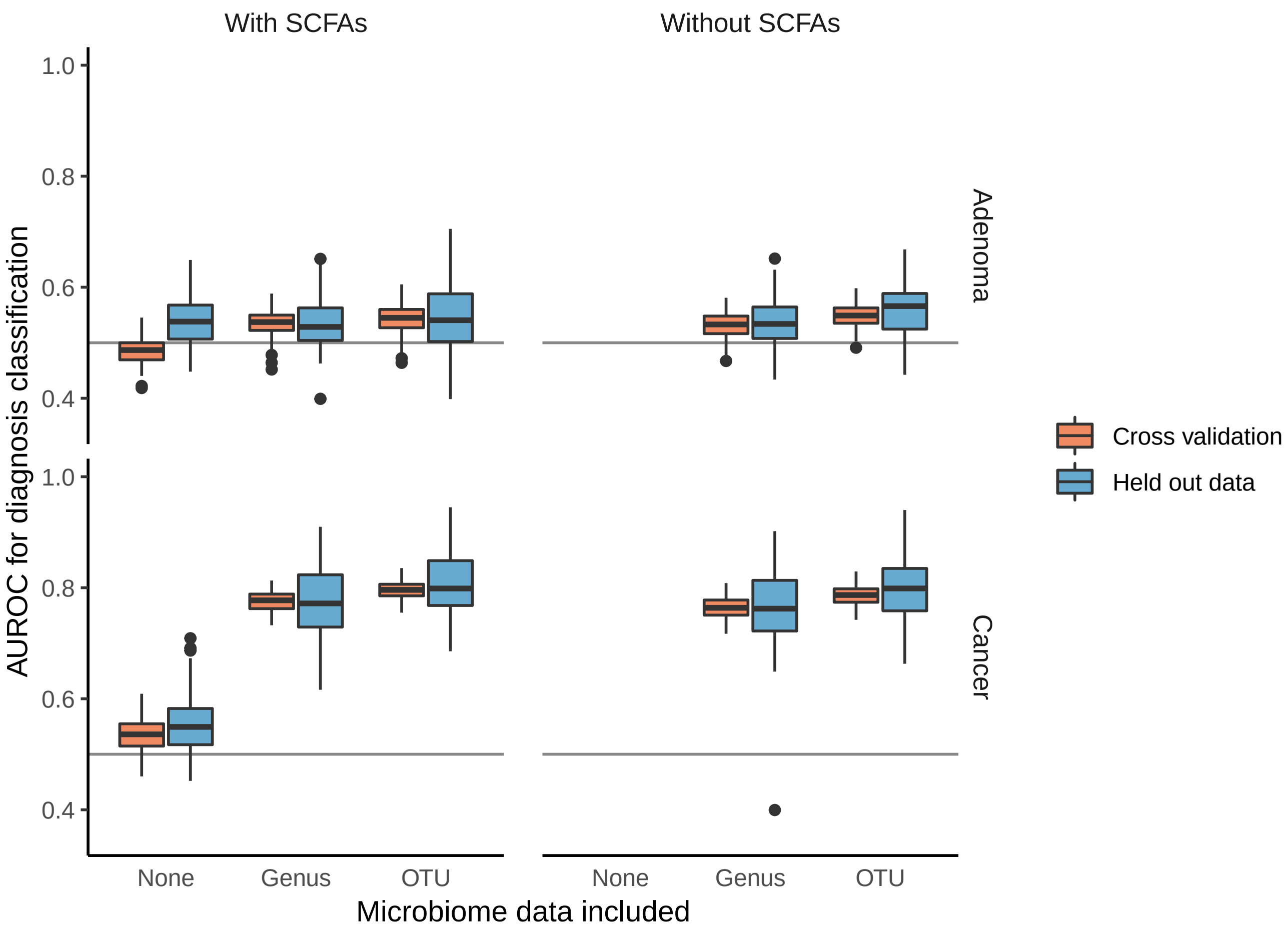

### Figure S2

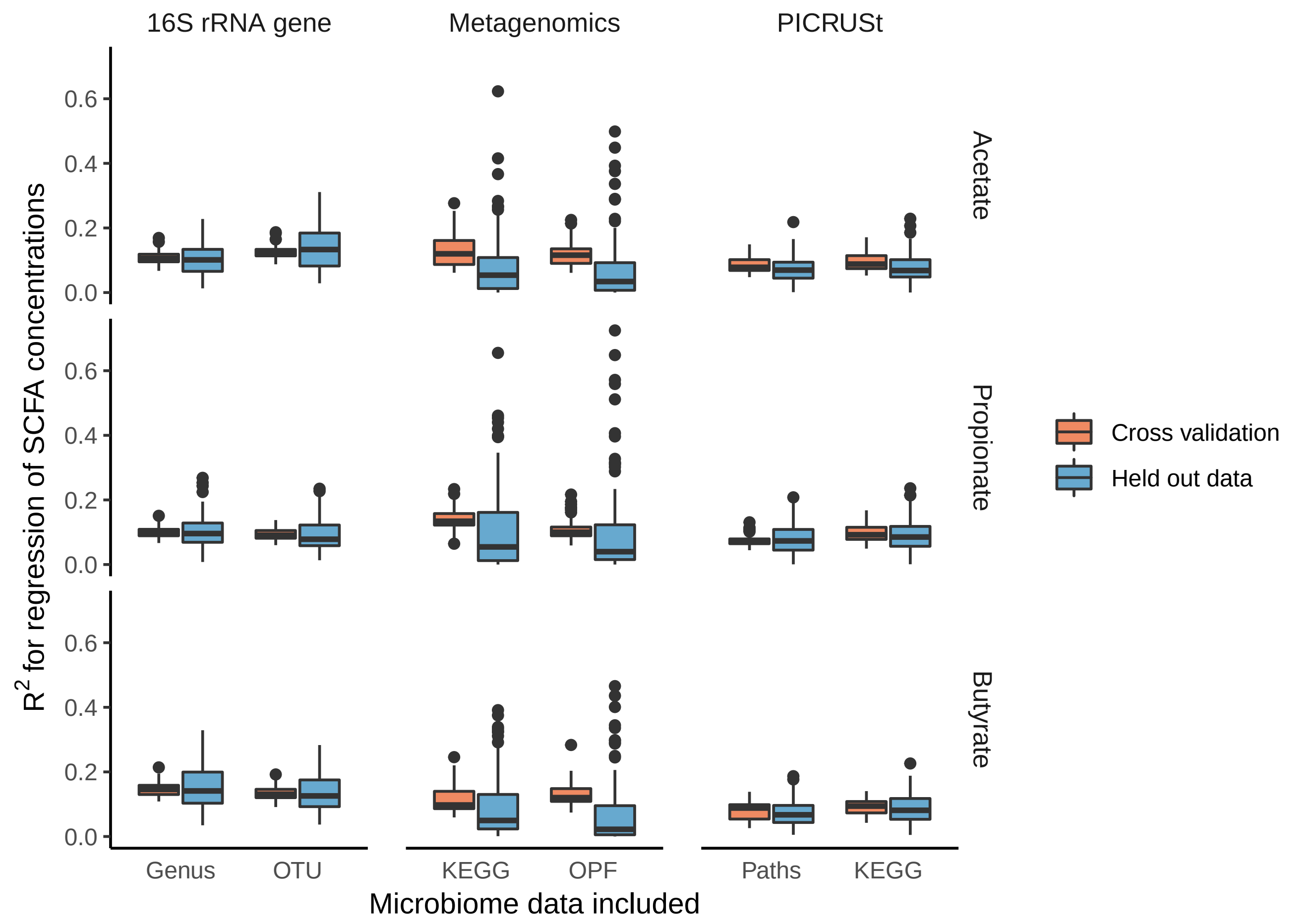
